# Advanced Cardiac Organoid Model for Studying Doxorubicin-Induced Cardiotoxicity

**DOI:** 10.1101/2025.05.02.651878

**Authors:** Xian Wu, Savanna Williams, Jacques Robidoux, Srinivas Sriramula, Abdel-Rahman Abdel

## Abstract

Cardiac organoids provide an in vitro platform for studying heart disease mechanisms and drug responses. However, a major limitation is the immaturity of cardiomyocytes, restricting their ability to mimic adult cardiac physiology. Additionally, the inadequacy of commonly used extracellular matrices (ECM), which fail to replicate the biochemical and mechanical properties of natural heart tissue, poses significant challenges. Consequently, structural integrity in cardiac organoids is impaired. Moreover, scalability remains an obstacle, as conventional ECM substitutes hinder mass production of organoids for high-throughput toxicology screening. To overcome these challenges, we developed an advanced model promoting fibroblast-driven ECM self-secretion, enabling physiologically relevant tissue architecture and function.

Using the ECM-free, mature cardiomyocyte-integrated organoid model, we investigated the cardiotoxicity of doxorubicin, a widely used chemotherapeutic agent known to impair cardiac function. Cardiomyocytes derived from induced pluripotent stem cells were characterized for maturity by immunostaining for cTNT and MYL2 alongside gene expression analysis. Organoids treated with doxorubicin showed reduced size and increased collagen deposition. These structural changes correlated with functional impairments, including decreased contraction rate and disrupted synchronous beating. In 2D culture, exposure to doxorubicin induced fibroblast activation, promoted endothelial-to-mesenchymal transition in endothelial cells, and triggered cytotoxic effects in cardiomyocytes.

This study highlights the importance of ECM remodeling in advancing cardiac organoid models and demonstrates its potential for more accurate cardiotoxicity assessment. Addressing these limitations enhances the physiological relevance of cardiac organoid systems for drug safety assessment and cardiac disease modeling.

## Introduction

Cardiac organoids are three-dimensional structures derived from stem cells replicating the cellular configuration and function of the human heart (Sahara 2023). They are small in-vitro models physiologically suited for cardiac disease mechanisms and drug responses compared to standard two-dimensional cultures. Organoids have spontaneous contractions, output in response to pharmaceuticals, and action potential patterns comparable to those of the human heart (Thomas et al. 2022). Cardiac organoids hold great significance in medical and toxicology research (Matsui and Shinozawa 2021). They provide a valuable platform for revealing mechanisms of cardiac diseases, drug screening, and understanding genetic mutations (Zhao, Lei, and Hu 2021). However, a key limitation is cardiomyocyte immaturity derived from human induced pluripotent stem cells (iPSCs), affecting their ability to recapitulate adult heart physiology and disease (Richards et al. 2020). Within cardiac organoids, one important component is the extracellular matrix, which serves as the structural and biochemical supporter for cells. Collagen I is a major fibrous protein in adult heart contributing to myocardial structure and function (Singh, Rai, and Agrawal 2023). Reconstructing ECM within cardiac organoids is a prerequisite for a precise model of cardiac physiology and pathology (de Castro Bras and Frangogiannis 2020). The currently used matrices, such as matrigel and hydrogels, have several disadvantages. Batch-to-batch variability in matrigel leads to unreliable results, and these materials do not accurately mimic the stiffness and elasticity of native human cardiac tissue (Kim et al. 2022). This discrepancy can influence cell behavior, differentiation, and the maturation of cardiomyocytes within the organoid. The ECM of the human heart is specialized; matrigel and generic hydrogels cannot replicate its intricate composition, limiting organoid physiological relevance (Min et al. 2024). Producing large quantities of organoids using matrigel or hydrogels is challenging due to handling difficulties, impeding scalability for toxicology screening. In contrast, ECM-free organoids adapted for high-throughput culture plates provide a practical and scalable solution for cardiotoxicity research. Therefore, we aim to enhance the organoid model by promoting the self-secretion of extracellular matrix components like collagen from fibroblasts to mimic the natural extracellular matrix of the heart.

Doxorubicin, as a highly effective chemotherapy drug has long been used in the treatment of cancers, such as leukemias, lymphomas, and solid tumors (Kciuk et al. 2023). It intercalates into DNA, which stops the replication of rapidly growing cancer cells (Rivankar 2014). Despite its use for cancer therapy, doxorubicin is well-known for its severe cardiotoxicity side effect (Kciuk et al. 2023). The dose-dependent toxicity of this drug primarily leads to arrhythmias, cardiomyopathy, and ultimately heart failure (Rawat et al. 2021). At the molecular level, Doxorubicin treatment results in an elevation of reactive oxygen species (ROS) in heart, resulting mitochondrial dysfunction and activation of apoptotic pathways in cardiac cells (Renu et al. 2018; Rawat et al. 2021). Doxorubicin cardiotoxicity also disturbs cardiac electrical activity causing arrhythmogenic effects like irregular heartbeats, further impairing heart function (Kilickap et al. 2007). These effects affect the structural integrity and proper functioning of heart muscle, further leading to progressive fibrosis and diminution of heart. The knowledge of the mechanisms underlying cardiotoxicity of doxorubicin is essential for developing strategies to prevent/reduce these negative effects and enhances quality of life of cancer patients.

Understanding fibrosis, electrophysiologic disturbances, and irregular beating patterns in organoids reflecting doxorubicin-induced arrhythmias is valuable for several reasons. Initially, cardiac organoids, particularly with matured cardiomyocytes, serve as better, more human-relevant platforms to investigate cardiotoxicities elicited by doxorubicin compared to conventional models such as animal studies or two-dimensional cell cultures. The organoid platform mimics the intricate cell-cell and cell-ECM interactions seen in the human heart, and matured cardiomyocytes are closer in functional nature to adult heart cells. Next, exploring the processes of fibrosis and arrhythmic contractions by doxorubicin in cardiac organoids could give new ideas on the early-stage cardiotoxicity which is complicated to indicate in clinic until severe damage occurs. It is significant that maturation of cardiomyocytes made the result closer to the adult heart. Furthermore, a molecular understanding of gene and protein disruption can help identify potential biomarkers for early detection of cardiotoxicity and guide the development of targeted therapeutics.

In this study, we aimed to create ECM-free matured cardiac organoids using iPSC-derived mature cardiomyocytes, cardiac fibroblasts, and endothelial cells to model the structure and function of human heart tissue. Additionally, we sought to validate the model by investigating how doxorubicin induces fibrosis and arrhythmic contractions through fibroblast activation, collagen I accumulation, endothelial-to-mesenchymal transition (EndMT), and cardiomyocyte cytotoxicity.

## Methods

### Stem cell maintenance and cardiomyocyte differentiation

Human induced pluripotent stem cells (iPSCs) (ND50019, Infinity Biologix), derived from human fibroblasts, were maintained in StemFlex medium (A3349401, Thermo Fisher) until reaching 80–90% confluence. Cells were passaged three times before differentiation. Cardiomyocyte differentiation was performed in 60-mm cell culture dishes (353219, Corning) using the monolayer differentiation method (Dark et al. 2023). Purification was conducted in glucose-free RPMI 1640 medium for 3 days before transitioning to maturation medium. Cardiomyocytes were then maintained in maturation medium following the protocol published (Feyen et al. 2020) for 40 days before following experimentation.

To generate cardiac organoids, 200 µL of cell suspension containing 150,000 iPSC-derived cardiomyocytes, 90,000 cardiac fibroblasts (CC-2904, Lonza), and 60,000 HUVECs (C2517A, Lonza) was pipetted into a 96-well ultra-low attachment plate. The plate was centrifuged at 1,000 rpm for 5 minutes and incubated overnight with 10 µM ROCK inhibitor Y-27632 (S1049, Selleckchem). On the second day, the medium was replaced without ROCK inhibitor, and medium changes were performed every 2 days throughout the doxorubicin exposure period. Cardiomyocytes were incubated in glucose-containing, lipid-rich maturation medium, prepared according to protocol in Feyen’s lab. Cardiac fibroblasts were cultured in fibroblast growth medium (CC-3132, Lonza), while HUVECs were maintained in endothelial cell growth medium (CC-3162, Lonza). Cardiac organoids were cultured in a mixed medium formulation with cardiomyocyte, fibroblast and endothelial cell medium according to the initial cell ratio. The experiment commenced on Day 0 (D0) after 10 days of organoid self-assembly. From D0 to D3, cardiac organoids were exposed to either the control or doxorubicin treatment protocol.

### Chemical exposure

Cardiac organoids were exposed to doxorubicin hydrochloride (AAJ64000MA, Fisher Scientific) at 0, 0.01, 0.1 and 1 µM for 3 days. Following treatment, organoids underwent a 7-day recovery period in doxorubicin-free medium before functional characterization and immunoassay analysis. To assess the effect of adrenergic stimulation, norepinephrine (A7257, Millipore Sigma) was added to cardiac organoids at final concentrations of 0, 0.25, 0.5 and 1 µM for 20 minutes prior to contraction rate analysis. For cell viability assays, monocultured cardiomyocytes, fibroblasts, and HUVECs were treated with doxorubicin (0, 0.001, 0.01, 0.1, 1 and 10 µM) for 1, 2 and 3 days. For immunoassay analysis, fibroblasts and HUVECs were treated with 0.1 µM doxorubicin for 3 days before further assessment.

### MTT viability assay

Cell viability was assessed using the MTT assay (3-(4,5-dimethylthiazol-2-yl)-2,5-diphenyl tetrazolium bromide). Cardiomyocytes, fibroblasts, and HUVECs were seeded in 96-well plates at a density of 10,000 cells/well. After 24 hours, the culture medium was replaced with fresh medium containing doxorubicin (0, 0.001, 0.01, 0.1, 1, or 10 µM), prepared in cell type-specific media. Cells were exposed to doxorubicin for 1, 2, or 3 days. At each time point, the treatment medium was replaced with 100 µL of fresh medium: cardiomyocytes in maturation medium, fibroblasts in fibroblast growth medium (CC-3132, Lonza), and HUVECs in endothelial cell growth medium (CC-3162, Lonza). Next, 10 µL of 12 mM MTT stock solution was added to each well, including medium-only negative controls. The plate was incubated at 37°C for 4 hours. Following incubation, 25 µL of supernatant was removed, and 50 µL of DMSO was added to each well. The plate was mixed thoroughly, incubated at 37°C for 10 minutes, mixed again, and absorbance was measured at 540 nm using a microplate reader.

### Video acquisition and analysis

The contraction of cardiac organoids, with and without chemical exposure, was recorded using a Celldiscoverer 7 (ZEISS, NY) maintained at 37°C and 5% CO□. Organoids were recorded after a total of 10 days, consisting of 3 days of doxorubicin exposure followed by 7 days of recovery. Videos were captured at 13 frames per second (fps), and contraction rates were quantified for 40 s per organoid by analyzing region-of-interest intensity changes using ZEN software (ZEISS, NY).

### Western blot

Attached cells were dissociated using TrypLE solution, followed by lysis in RIPA buffer (20-188; EMD Millipore) containing protease inhibitor cocktail. Total protein concentrations were determined using a bicinchoninic acid (BCA) assay kit (23225; Thermo Fisher Scientific). Samples (20 µg protein/sample) were separated by sodium dodecyl sulfate–polyacrylamide gel electrophoresis (SDS-PAGE) on 4%–12% bis-Tris gels (3450124; Bio-Rad). Proteins were transferred onto 0.45-µm nitrocellulose membranes (1620115; Bio-Rad) and blocked in PBS-based blocking solution [1× PBS, 0.1% Tween-20, and 3% (w/v) bovine serum albumin (BSA)] for 1 h at room temperature. Membranes were incubated overnight at 4°C with primary antibodies: cardiac troponin T (MA5-12960; Thermo Fisher Scientific, 1:200), MYL2 (ab79935; Abcam, 1:200), COL1A1 (72026; Cell Signaling, 1:200), vimentin (5741; Cell Signaling, 1:200), CD31 (3528; Cell Signaling, 1:200), and α-smooth muscle actin (19245; Cell Signaling, 1:200). After washing, membranes were incubated with secondary antibodies (1:1000 dilution in blocking buffer) for 1 h at room temperature with gentle agitation. Protein bands were visualized using the Odyssey imaging system (LI-COR) and quantified using Image Studio software (version 5.0).

### Realtime PCR

RNA was extracted using the RNeasy kit (R2050; ZYMO Research), and its quality were assessed spectrophotometrically based on the 260/280 nm absorbance ratio. Complementary DNA (cDNA) was synthesized from 500 ng of total RNA using the QuantiTect Reverse Transcription kit. Quantitative PCR (qPCR) was performed using 20 ng of cDNA per reaction, gene-specific primers, and Fast SYBR™ Green Master Mix. Thermal cycling conditions were as follows: initial denaturation at 95°C for 5 min, followed by 40 cycles of 95°C for 20 s, 60°C for 20 s, and 72°C for 20 s, and a final extension step at 72°C for 5 min. Melt-curve analysis was conducted to confirm the specificity of PCR products. Reactions were run on a QuantStudio™ 7 Flex Real-Time PCR system (Applied Biosystems), and data were analyzed using QuantStudio™ software. Target gene expression levels were normalized to the GAPDH housekeeping gene and calculated using the 2^–ΔΔCt method.

### Statistical Analysis

Statistical analyses in the figure generation were performed using Prism software (version 10.0.2; GraphPad Software). Data comparisons among independent groups were analyzed by one-way analysis of variance (ANOVA), followed by appropriate post-hoc tests to determine significant differences. Differences in gene expression between two groups were analyzed using a two-tailed *t*-test. Median lethal concentration (LC□□) values were determined using nonlinear regression analysis in Prism. Statistical significance was defined as *p* < 0.05 for all analyses.

## Results

### Cardiomyocytes derived from iPSCs mature in presence of maturation medium

Human iPSCs differentiated into cardiomyocytes by regulating WNT signaling and started beating from day 8, mimicking the early stage of cardiac cell formation(Wu et al. 2022). As cardiomyocytes mature following iPSC differentiation, they undergo a metabolic shift from primarily utilizing glycolysis to fatty acid oxidation and begin expressing maturation marker proteins. Immunostaining of day 40 (D40) cardiomyocytes in maturation medium revealed positive markers for cTNT and MYL2 (markers for mature ventricular cardiomyocytes), confirming their maturation (Fig. 1A). To further characterize this maturation process at the gene expression level, RNA was extracted from iPSCs, day 10 (D10) cardiomyocytes in maintenance medium, and D40 cardiomyocytes in maturation medium, and analyzed via real-time PCR for mRNA expression. Notably, the expression of *CPT1B* and *SLC25A20* (markers for fatty acid oxidation) significantly increased in D40 cardiomyocytes compared to the immature D10 cells (Fig. 1B). The cardiac potassium ion channel *KCNJ2* and sodium channel *SCN5A* also showed significant increases in D40 cardiomyocytes, reaching levels comparable to those in the human heart. The maturation-specific markers such as MYL2, *CPT1B, SLC25A20, KCNJ2* and *SCN5A* distinguish mature cardiomyocytes from their immature counterparts, highlight the metabolic and functional maturation of iPSC-derived cardiomyocytes in presence of maturation medium.

**Figure 1.**
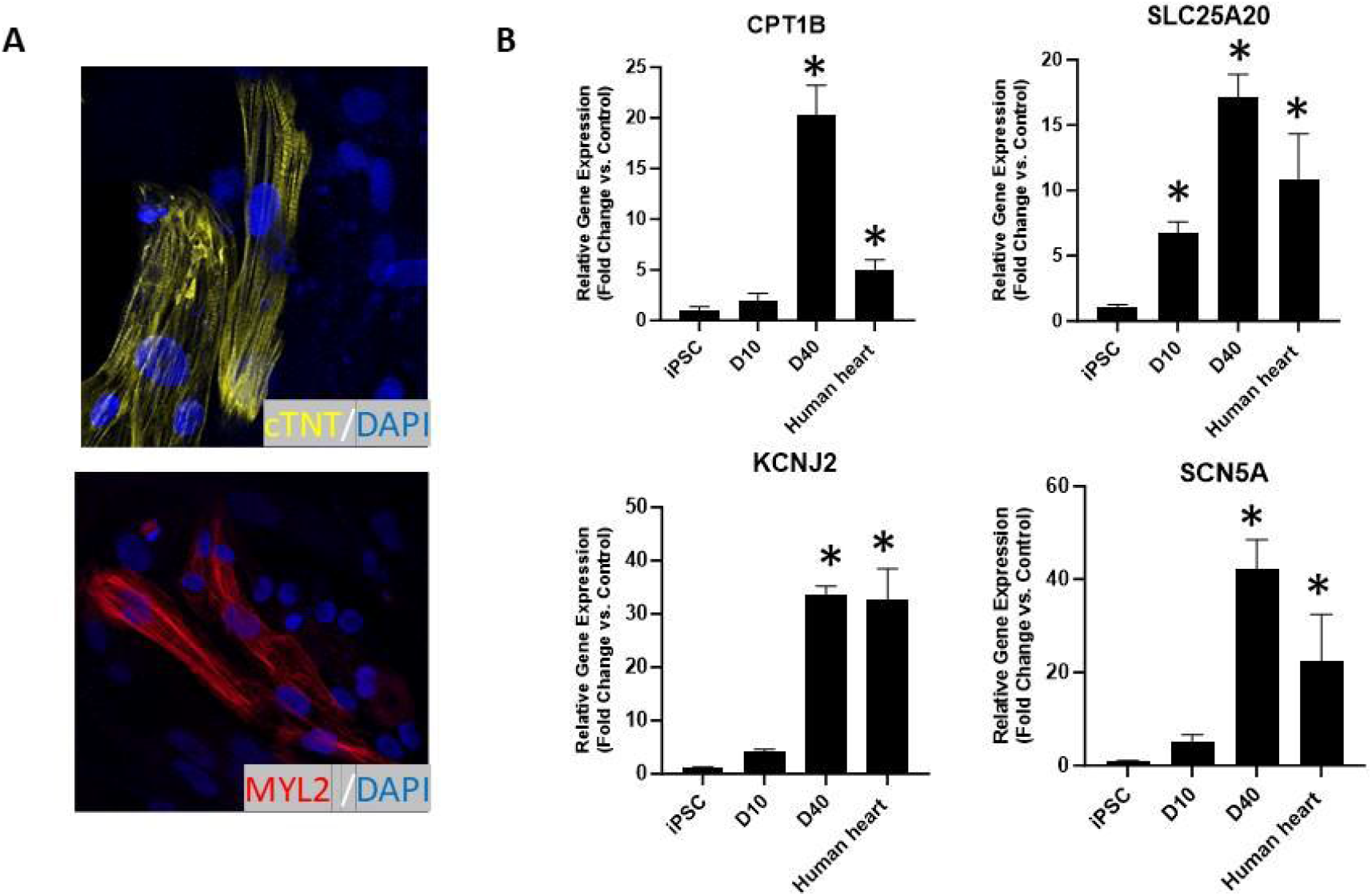
Characterization of iPSC-derived mature cardiomyocytes. (A) Immunofluorescence staining of human iPSC-derived cardiomyocytes using mature cardiomyocyte-specific antibodies (cTnT, MYL2). (B) Gene expression analysis by RT-PCR comparing selected mature cardiomyocyte markers at day 10 (D10) and day 40 (D40). Data presented as mean fold-change (2−ΔΔCt) ± SD (*p < 0.05; n = 3, with 10 cardiac organoids per group). CPT1B: carnitine palmitoyltransferase 1B; SLC25A20: mitochondrial carnitine/acylcarnitine carrier protein; KCNJ2: potassium inwardly rectifying channel subfamily J member 2; SCN5A: sodium voltage-gated channel alpha subunit 5.

### Norepinephrine increases self-aggregated cardiac organoids contractility

To engineer and validate multicellular cardiac organoid for drug toxicity testing, next we looked at organoids composite of mature cardiomyocytes, along with endothelial cells and fibroblasts, aggregated to form cardiac organoids within 10 days (Fig. 2A). To confirm the cell type involved, these organoids were fixed in 4% paraformaldehyde, and immunostaining revealed that fibroblasts (vimentin), endothelial cells (CD31), and cardiomyocytes (MYL2) were distributed throughout the organoid (Fig. 2A). After 10 days, the cardiac organoids secreted collagen (COL1A1) and responded to norepinephrine exposure indicating self-aggregation and physiological responsiveness of the cardiac organoid. Specifically, exposure to 0.5 and 1 µM norepinephrine significantly increased the beating frequency of the cardiac organoids compared to the untreated group (Fig. 2B) demonstrating functional adrenergic receptor activity and the dose dependent responsiveness of the organoids to drug stimulation.

**Figure 2.**
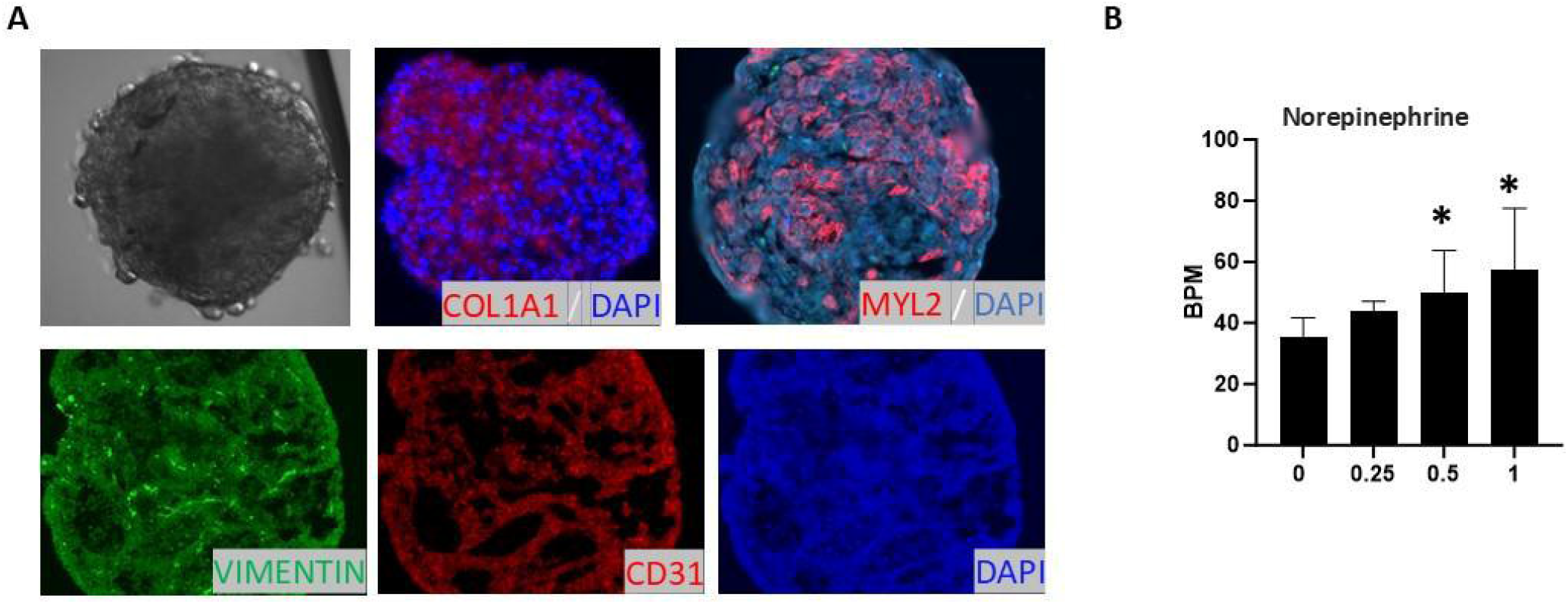
Cardiac organoid formation and characterization. (A) Self-organized cardiac organoids immunostained for fibroblasts (vimentin), endothelial cells (CD31), cardiomyocytes (MYL2), and extracellular matrix component collagen I (COL1A1). (B) Contractility assessment of cardiac organoids at day 10 (D10). Organoids were treated with norepinephrine (0.25, 0.5, and 1 µM) for 20 min, and beating rates (beats per minute, BPM) were quantified through video-based region-of-interest (ROI) analysis (*p < 0.05; n = 10).

### Doxorubicin disrupts the cardiac organoid formation and contractility

Next, we investigated the effects of doxorubicin using the validated organoid model to mimics the doxorubicin effect in adult heart. The morphology and function of cardiac organoids were assessed after 3 days of doxorubicin exposure followed by a 7-day recovery period. Doxorubicin treatment reduced organoid diameter to 376 µm at a concentration of 0.01 µM and to 365 µm at 0.1 µM, compared to the average size of untreated organoids, which was 393 µm (Fig. 3A), indicating the dose dependent cytotoxic effect of doxorubicin in the organoid model. While 0.01 µM doxorubicin did not significantly affect organoid contractility, 0.1 µM doxorubicin reduced the beating rate from 49 BPM to 20 BPM (Fig. 3B) and disrupted rhythm (Fig. 3C). At 1 µM, organoids ceased beating entirely suggesting severe cytotoxicity.

**Figure 3.**
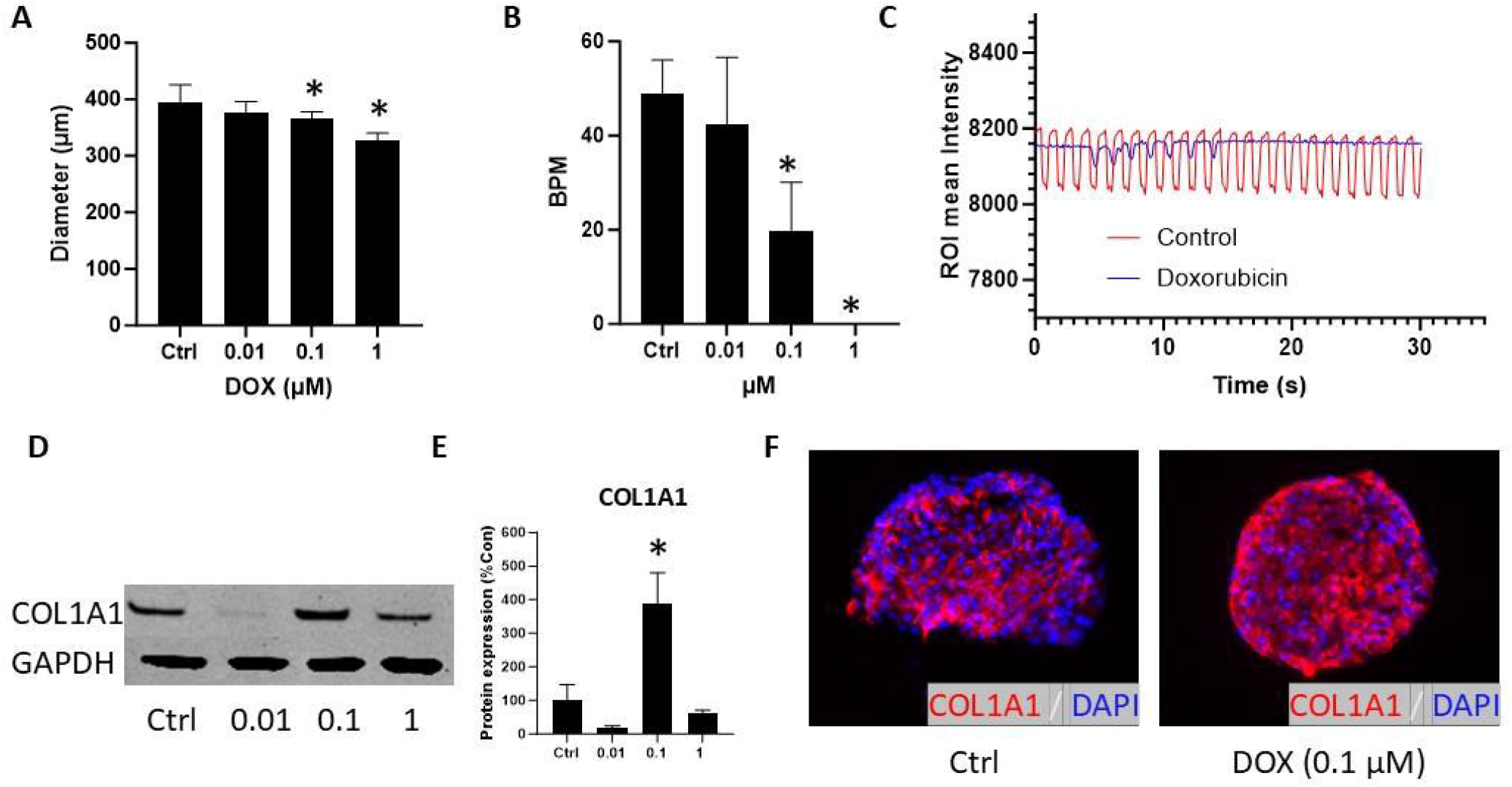
Doxorubicin disrupts cardiac organoid contractility. Cardiac organoids were cultured for 10 days in a 96-well ultralow attachment plate before exposure to doxorubicin for 3 days, followed by a 7-day recovery period. (A) Organoid size quantified by ImageJ analysis (*p < 0.05; n=10). (B) Contractility measured by video analysis; beating rates (BPM) were quantified (*p < 0.05; n=10). (C) Rhythm visualization of representative contraction patterns. (D, E) Western blot analysis of collagen I (COL1A1) protein expression and quantification (*p < 0.05; n=3, 10 organoids per group). (F) Immunofluorescence staining for COL1A1.

To further investigate the effects of doxorubicin on collagen I expression indicate the ECM integrity, COL1A1 levels were analyzed by western blot (Fig. 3D). Quantification showed no increase in COL1A1 expression in the 0.01 µM group, but a significant increase at 0.1 µM. No change was observed in the 1 µM group (Fig. 3E). Immunostaining confirmed elevated COL1A1 protein levels in the 0.1 µM group (Fig. 3F).

### Doxorubicin activates cardiac fibroblast cells

To further investigate the effects of doxorubicin on independent specific cell subtypes within the organoid, fibroblasts were cultured in 96-well plates for a viability analysis over 3 days. On day 1, doxorubicin did not affect fibroblast viability until the concentration reached 10 µM. By day 2 and day 3, doxorubicin significantly inhibited fibroblast viability at 1 µM and 10 µM (Fig. 4A). After 3 days of doxorubicin exposure, alpha smooth muscle expression in fibroblast cells increased notably at non cytotoxic 0.1 µM at the protein level (Fig. 4B, C).

**Figure 4.**
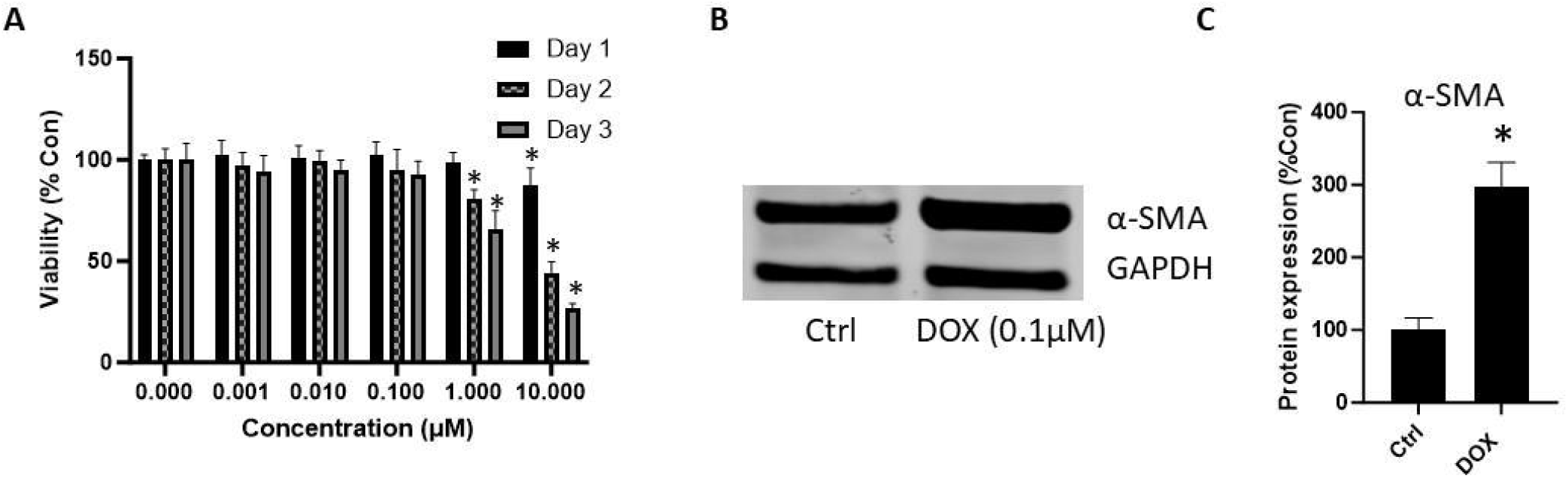
Doxorubicin-induced activation of cardiac fibroblasts. (A) Viability of fibroblasts seeded at 10,000 cells/well assessed by MTT assay following exposure to doxorubicin (0.1 µM) for 1, 2, and 3 days (*p < 0.05; n=8). (B, C) Western blot analysis and quantification of α-smooth muscle actin (α-SMA) expression in fibroblasts exposed to 0.1 µM doxorubicin.

### Doxorubicin induces HUVEC endothelial-to-mesenchymal transition

In the HUVEC endothelial cells, doxorubicin inhibited cell viability at 10 µM on day 1, and at 1 µM and 10 µM on day 2. By day 3, HUVEC cell viability decreased at concentrations as low as µM, 1 µM, and 10 µM (Fig. 5C) indicating that endothelial cells were susceptible to doxorubicin induced cytotoxicity during prolonged exposure. Additionally, after 3 days of exposure to 0.1 µM doxorubicin, HUVEC exhibited increased expression of alpha smooth muscle actin while the endothelial marker CD31 was downregulated, indicating endothelial-to-mesenchymal transition induced by doxorubicin (Fig. 5A, B, D).

**Figure 5.**
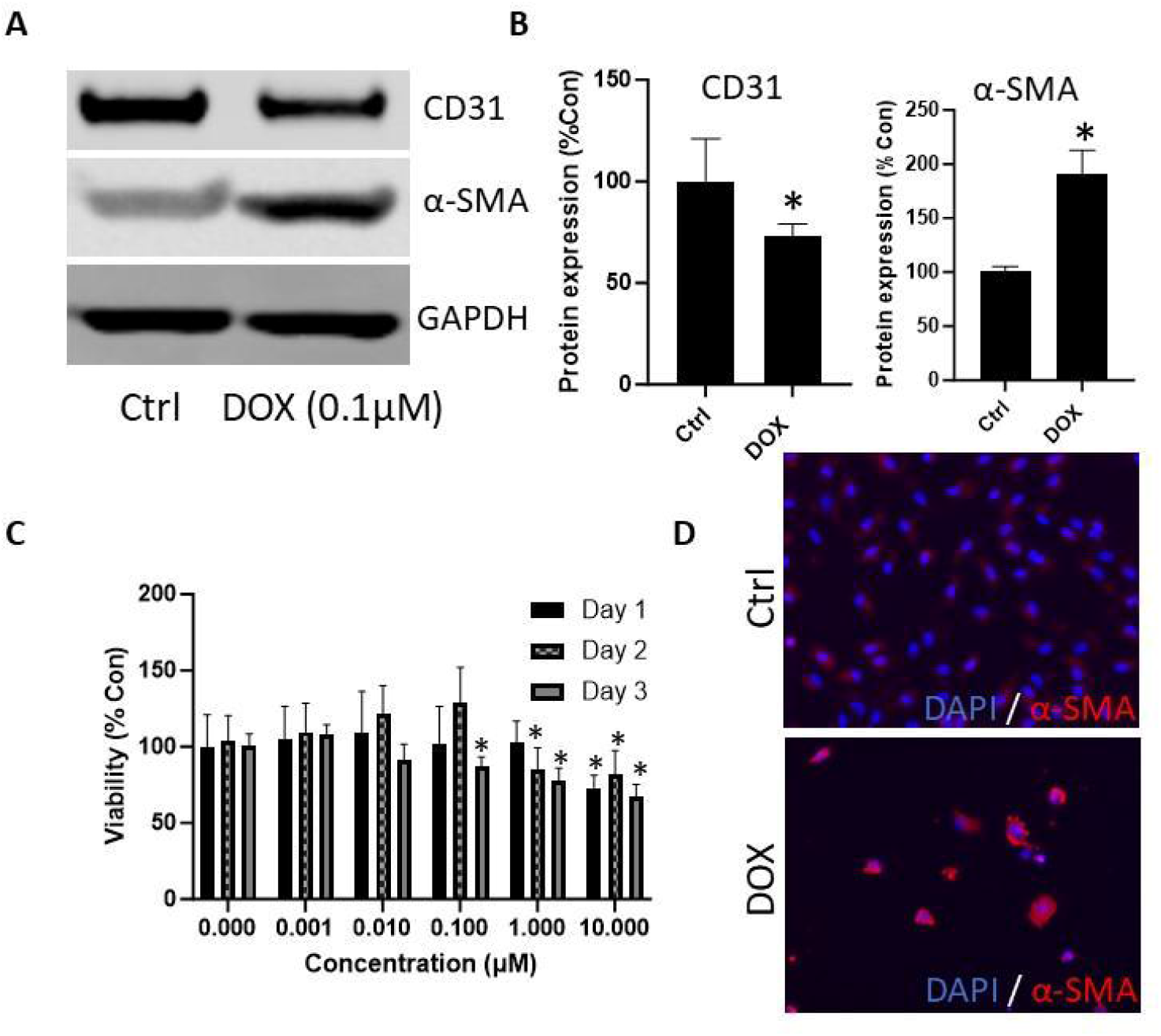
Doxorubicin induces endothelial cell toxicity and endothelial-to-mesenchymal transition. Endothelial cells were seeded at 10,000 cells/well, exposed to doxorubicin (0, 0.01, 0.1, 1 µM), and assessed for viability by MTT assay at 1, 2 and 3 days post-treatment (*p < 0.05; n=8). (A, B) Western blot analysis and quantification of α-SMA protein expression after 0.1 µM doxorubicin exposure (*p < 0.05; n=3, 10 organoids per group). (C) MTT assay evaluating cell viability after doxorubicin treatment on days 1–3. (D) Representative immunofluorescence staining demonstrating doxorubicin-induced α-SMA changes in endothelial cells.

### Doxorubicin reduces cardiomyocyte cell viability

Cardiomyocytes are the primary functional cells responsible for organoid contractility. To further investigate the effects of doxorubicin-induced stress on cardiomyocytes, mature cardiomyocytes derived from iPSC differentiation were exposed to doxorubicin for 3 days. After exposure, cardiomyocyte viability decreased significantly in a dose-dependent manner at concentrations ranging from 0.01 µM to 1 µM, highlighting the cytotoxic effects and the high sensitivity of cardiomyocytes to doxorubicin exposure (Fig. 6).

**Figure 6.**
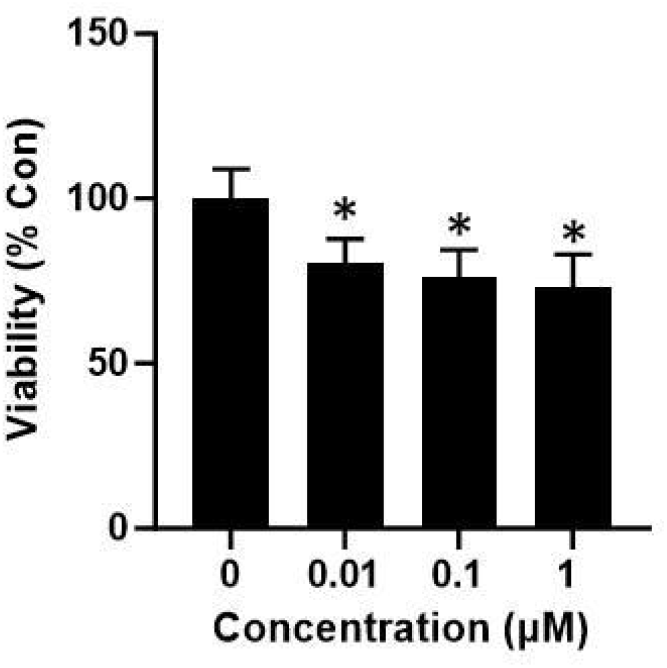
Doxorubicin reduces cardiomyocyte viability. Mature cardiomyocytes were seeded in 96-well plates at a density of 10,000 cells per well. Cells were exposed to doxorubicin (0, 0.01, 0.1, 1 µM) starting one day post-seeding, and viability was measured by MTT assay after 3 days (*p < 0.05; n=8).

## Discussion

The hiPSC-CMs hold great promise for studying human cardiac disorders, but their physiological immaturity limits their use in drug discovery (Li et al. 2024). Maturation media used in this study for cardiac organoid formation tailored to their metabolic needs enhances hiPSC-CMs function and maturation, The significant increase in the expression of *CPT1B* and *SLC25A20* after 40 days of differentiation, along with the positive immunostaining for cTNT and MYL2, confirms the maturation of the cardiomyocytes. Carnitine palmitoyltransferase 1B (*CPT1B*), a key regulator of FAO, plays a critical role in cardiomyocyte maturation. *SLC25A20* is a mitochondrial carnitine/acylcarnitine translocase, plays a critical role in fatty acid oxidation by transporting acylcarnitines into the mitochondrial matrix for β-oxidation. Evaluating *SLC25A20* alongside other maturation markers provides a comprehensive understanding of cardiomyocyte maturation. *CPT1B* and *SLC25A20* work together to facilitate the efficient transport and oxidation of long-chain fatty acids in adult cardiomyocytes, which is essential for meeting the high energy demands of the heart (Duan, Liu, and Zhan 2022). Their coordinated interaction ensures that the heart muscle can produce ATP through the oxidation of fatty acids to maintain its function and health. *KCNJ2* encodes the inwardly rectifying potassium channel Kir2.1, which is an important regulator of cardiac resting membrane potential and contributes to the cardiac action potential(Ponce-Balbuena et al. 2018). *KCNJ2* is essential for the maturation of cardiomyocytes, maintaining cardiac excitability and ensuring effective cardiac function. *SCN5A* encodes the voltage-gated sodium channel NaV1.5, which plays a central role in the initiation and propagation of the cardiac action potential. The generation and propagation of action potentials is the basis for normal cardiac rhythm and contractile synchronization (Abriel and Kass 2005). This 2D cardiomyocyte model has been reliably used in published studies to model genetic cardiac diseases: long QT syndrome and dilated cardiomyopathy, as well as hypoxia-induced cardiac damage (Feyen et al. 2020). In addition, 3D cardiac organoid incorporates matured cardiomyocyte provide a physiologically relevant 3D model that mimics the function of the human heart, enabling studies of heart disease mechanisms. The self-organization of cardiac fibroblasts, endothelial cells and mature cardiomyocytes in this study provides a functional and simplified in vitro model for high-throughput screening in pharmacological and toxicological research compared to existing cardiac organoids in the literature (Zhao, Lei, and Hu 2021). Their ability to secrete collagen I and respond to norepinephrine exposure, underscores their functional relevance to adult human heart. The significant increase in beating frequency upon norepinephrine exposure highlights their physiological responsiveness. Currently, cardiac organoids have used extensive additional ECM, either from rodents (Matrigel) or engineered hydrogels, which have several disadvantages including batch-to-batch variability which will not accurately mimic the stiffness and elasticity of native human cardiac tissue (Kaur et al. 2021). This discrepancy can influence cell behavior, differentiation, and the maturation of cardiomyocytes within the organoid. The ECM of the human heart is highly specialized, and Matrigel and generic hydrogels cannot fully replicate its intricate composition and organization, limiting the physiological relevance of the organoids. We have validated in this study that cardiac organoids that self-organize *in vitro* without additional ECM are functional with contractility function similar to human heart and showed that doxorubicin disrupts the cardiac organoid contractility (Figure. 3).

To validate the advanced cardiac organoid model, the effects of the chemotherapy drug doxorubicin on cardiac organoids were assessed to replicate the heart’s response within this model system. In cancer patients, doxorubicin blood plasma levels range between 0.02 and 1.14 μM (Harahap et al. 2020) and persist for up to 72 hours (Jackson et al. 2019). Based on this and other literature (Fant et al. 2019), a 72-hour exposure to doxorubicin at doses of 0.01, 0.1, and 1 μM, followed by a 7-day recovery, was designed to mimic the first cycle of doxorubicin therapy in patients. This model demonstrated doxorubicin reduced the contractility of cardiac organoids at 0.1 µM and completely halted beating at 1 µM after 3 days of exposure followed by a 7-day recovery period. The untreated cardiac organoids maintained a consistent beating frequency, which was significantly reduced in the 0.1 µM treated group, with an increase in beating intervals by 15 seconds. The disturbed beating frequency in the doxorubicin 0.1 µM treated group also demonstrates the doxorubicin’s arrhythmogenic effect (Figure. 3C). Our findings are consistent with previous studies that have documented the cardiotoxic effects of doxorubicin, including reduced contractility and induced arrhythmias in patient with doxorubicin chemotherapy (Benjanuwattra et al. 2020). Additionally, doxorubicin exposure led to a decrease in organoid size and an increase in alpha smooth muscle protein expression, indicating cellular stress and remodeling.

In the 2D cell model systems, doxorubicin activates cardiac fibroblast cells through upregulation of α-SMA expression and induces endothelial-to-mesenchymal transition (EndMT) in HUVECs, potentially contributing to the cardiac fibrosis that leads to the arrhythmic effects of doxorubicin. Previous literature has also reported doxorubicin-induced fibrosis and endothelial-to-mesenchymal transition (EndMT) (Tsai et al. 2019), which aligns with our observation of increased α-SMA expression in both fibroblasts and endothelial cells after doxorubicin exposure. The long-term extracellular matrix secretion from fibroblast in the 3D cardiac organoid model and mechanisms need to be further investigated.

This study has several limitations. The in vitro nature of cardiac organoids, while providing valuable insights, may not fully replicate the complex in vivo environment of the human heart particularly due to the absence of immune cell interactions and endocrine influence on the organoid system. Additionally, the iPSC-derived cardiomyocytes were not specifically differentiated to mimic ventricle or atrium, which may not capture whether the drug has specific adverse effects on different cardiac regions. Finally, incorporating more advanced techniques, such as single-cell RNA sequencing, could provide deeper insights into model characterization and the cellular and molecular changes induced by doxorubicin.

In closing, this project developed an ECM free self-organized and functionally matured 3D human cardiac organoid system, incorporating hiPSC-derived matured cardiomyocytes, fibroblast cells, and endothelial cells. This model addresses the common limitation of iPSC derived cardiac organoid, particularly cardiomyocytes immaturity in structural and functional properties, offering a more physiologically relevant system for cardiovascular research. The engineered cardiac organoid in 96 well plate format combined with imaging analysis in the study enables fast screening of doxorubicin cardiotoxicity in this research. Furthermore, this 3D cardiac organoid with 2D cellular models could be a valuable tool for other drugs cardiac toxicity screening and underlying mechanism exploration.

## Acknowledgements

The authors would like to thank research technician Mina Zhang in Dr. Abdel-Rahman’s lab at the East Carolina University, who assisted the cell culture and western blot, organoid cryosectioning.

